# Response of soil microbiome composition to selection on methane oxidation rate

**DOI:** 10.1101/2023.06.23.546315

**Authors:** Andrew H. Morris, Brendan J. M. Bohannan

## Abstract

Microbiomes mediate important ecosystem functions, yet it has proven difficult to determine the relationship between microbiome composition and the rate of ecosystem functions. This challenge remains because it is difficult to manipulate microbiome composition directly, we often cannot know *a priori* which microbiome members influence the rate of an ecosystem function, and microbiomes can covary strongly with other drivers of ecosystem function, such as the environment. To address these challenges, we imposed artificial selection on whole soil ecosystems over multiple generations to select for microbial communities with a high rate of CH_4_ oxidation. This approach is potentially powerful because it is biologically “agnostic” in that it makes few assumptions about which taxa are important to function, and repeated passaging with fresh substrate weakens the covariance between microbes and the environment. As a response to selection, we observed a 50.7% increase in CH_4_ oxidation rate per passage relative to a control that experienced random selection. We estimated that 31.5% of the variation in CH_4_ oxidation rate in these soils can be attributed to microbiome variation (though this was not significant). We also found that selection did not enrich for known CH_4_ oxidizers; instead, 12 families not known to oxidize CH_4_, including *Fimbriimonadaceae*, *Cytophagaceae*, and *Diplorickettsiaceae*, were enriched by selection. This result is in contrast to the typical assumption that the rate of an ecosystem function is limited by the final step in the associated microbial pathway. Our study demonstrates that variation in microbiome composition can contribute to variation in the rate of ecosystem function independent of the environment and that this may not always be limited by the final step in a pathway. This suggests that manipulating microbiome composition directly without altering the environment could be a viable strategy for managing ecosystem functions.

## Introduction

Microbiomes mediate a variety of important ecosystem functions relevant to human health, agriculture, and global change. As a result, there is great interest in understanding how to manipulate the microbiome to achieve desirable outcomes within these domains (1–3). However, for microbiome manipulations to be successful, variation in the microbiome must contribute directly to variation in the magnitude of the function of interest independent of other factors. Many studies have attempted to document such a relationship (4–9). However, it is difficult to isolate the direct effect of variation in microbiome composition from other drivers of variation in ecosystem function, such as the indirect effect of the environment on function through microbiome assembly. Here, we overcome these limitations by using a selection approach to estimate the degree to which an ecosystem function varies with microbiome composition.

Altering ecosystem functions via microbiome manipulations requires that the microbiome contributes to variation in ecosystem function independent of other drivers of ecosystem variation, such as variation in environmental conditions. This is because the drivers of variation in ecosystem functions can interact in complicated ways (Figure 1). Variation in the microbiome can contribute directly to variation in ecosystem function, for example, if a microbial population is replaced by one with a greater enzyme efficiency. In addition, environmental conditions can contribute indirectly to ecosystem function via covariance with the microbiome, for example, by providing conditions that select for microbial groups that in turn alter the rates of ecosystem functions. In this scenario, identifying the change in microbial community composition without adequately controlling for the environmental conditions would incorrectly attribute the change in ecosystem function to the microbiome when it is ultimately an indirect effect of the environment. Determining the independent contribution of microbiome variation to ecosystem function is crucial because if microbiome composition is driven primarily by environmental conditions, then introducing a desirable taxon through microbiome manipulation without altering the environment will likely be unsuccessful at shifting the targeted ecosystem function.

**Figure 1:**
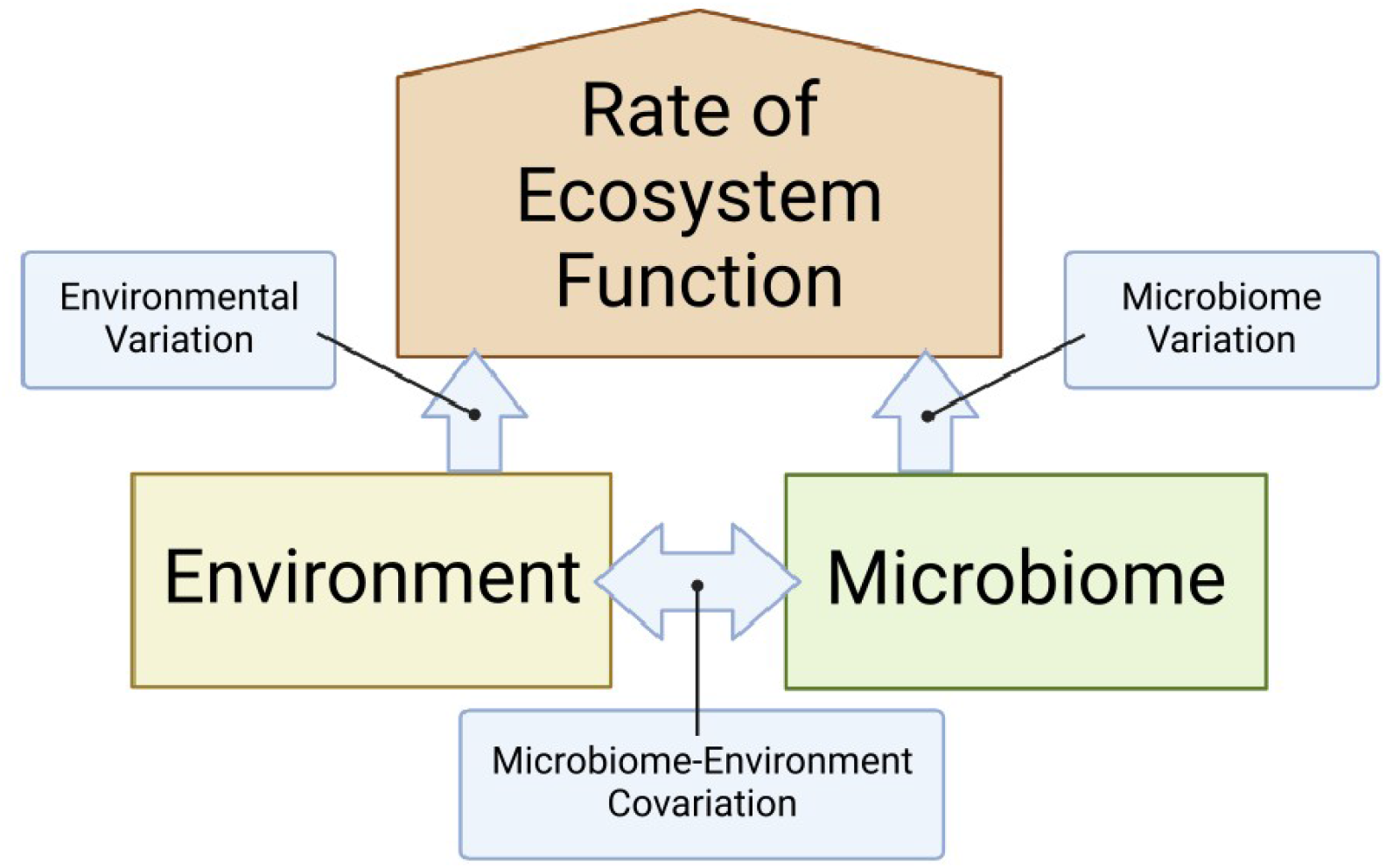
Variation in the rate of an ecosystem function is the result of at least three components: variation in the abiotic environmental conditions, variation in microbiome composition, and the covariance between microbiomes and the environment. The arrows represent causal relationships between the components. It is important to isolate the direct effect of the microbiome from the effect of the environment via covariance with the microbiome. Here, we attempt to isolate the effect of the microbiome through artificial selection on microbiome composition. For simplicity, we omitted the reverse arrows as well as the interactions, though these relationships may also exist.

There have been two general categories of approaches that investigators have used to estimate the degree to which an ecosystem function varies with microbiome composition: comparative and manipulative. Comparative studies sample natural variation in an ecosystem function across different habitats and simultaneously measure variation in community composition. Investigators can then correlate ecosystem function with aspects of community composition while attempting to control for environmental variation. These approaches have documented important relationships between microbiomes and ecosystem functions. For example, a meta-analysis of these studies observed a small but significant contribution of the microbiome to variation in ecosystem function after controlling for environmental variation (10). In addition, studies focusing on the correlation between the rate of an ecosystem function and the abundance of an associated marker gene (i.e., a gene that codes for a protein assumed to be involved in the ecosystem function) sometimes observe a significant correlation, though this relationship is rare and contingent upon both the function and the ecosystem sampled (11). However, comparative studies come with unique challenges and limitations. One issue is that microbiome attributes tend to covary with the abiotic conditions within an environment, and it is difficult to control for these abiotic variables in order to identify the unique contribution of the microbiome to ecosystem function. In addition, it is difficult to know *a priori* which environmental variables or community attributes to measure. Finally, while these approaches can establish a potential magnitude and direction for these relationships, it is often difficult to identify the taxa or genes that explain the connection between composition and function.

The other broad category of approaches used to address this question are manipulative approaches. Manipulative experiments try to alter microbial community composition and observe the effect on function. For example, reciprocal transplant and common garden experiments have shown that microbiomes originating from different ecosystems inoculated into the same substrate or introduced into a common environment display distinct functional rates (4–7). In addition, manipulating diversity by filtering communities by cell size or through dilution has been shown to alter the rate of ecosystem functions (8,9,12). However, manipulating the microbiome directly is challenging, and manipulative approaches often confound community composition with other factors. For example, reciprocal transplant and common garden experiments can confound community composition with the abiotic conditions introduced with the inoculum, while manipulating composition through dilution may confound composition with biomass (13).

In this study, we sought to build on the observations of comparative and manipulative studies by applying a different approach to the question of whether microbiome variation contributes to variation in the rate of an ecosystem function. We used artificial ecosystem selection to select for microbiomes that performed a greater rate of ecosystem function (14–16). We then tested whether variation in the microbiome contributed to variation in the rate of ecosystem function and identified microbiome attributes that might explain this relationship. There are several potential advantages to this approach for documenting the direct contribution of the microbiome to variation in ecosystem function and for investigating the mechanisms underlying those relationships. Through repeated passaging of microbiomes in a common environment, we can weaken the covariance between microbes and the environment by repeatedly diluting the influence of variation in abiotic conditions. In addition, our approach eliminates the need to generate microbiome variation through methods that are confounded with biomass or cell size. Lastly, by comparing our artificially selected community to a control community resulting from random selection, we can both control for changes in the environment over time and identify genes or taxa that are associated with the ecosystem function under selection.

We applied artificial ecosystem selection to soil microbiomes by selecting on soil methane (CH_4_) oxidation rate. We chose this function because CH_4_ is a globally important greenhouse gas and CH_4_ oxidation by soil bacteria is the primary biological sink for atmospheric CH_4_ (17). In addition, there is evidence that soil CH_4_ oxidation rate may vary with microbiome composition based on comparative studies in a variety of arctic and tropical ecosystems (18–21) as well as studies that manipulate methanotroph richness (22). Finally, methanotrophy is one of the most deeply conserved microbial physiologies and is represented in a narrow range of taxa, which suggests that the taxonomic composition of the microbiome is more likely to be associated with the rate of CH_4_ oxidation than other broader or more shallowly conserved functions (2,23).

In this study, we used artificial ecosystem selection on CH_4_ oxidation rate to address the following questions: Does variation in the relative abundance of microbial taxa contribute to variation in soil CH_4_ oxidation rate independent of the environment in our system? Which attributes of the microbiome are associated with variation in CH_4_ oxidation rate, and do these attributes match our assumptions about the factors that regulate CH_4_ oxidation rate in nature?

## Materials and Methods

### Experimental design

We performed an artificial ecosystem selection experiment (*sensu* (14)) by passaging replicate soil microbiomes. The trait we selected on was soil CH_4_ oxidation rate. Soil microcosms were incubated at room temperature in sealed 500 mL glass jars with a rubber septum for gas sampling. Each jar was sterilized with 70% ethanol and was composed of 45 g of autoclaved artificial potting mix, 5 g of living soil inoculum, and 3.5 mL of sterile deionized water to bring the soil to 60% of field capacity. The potting mix consisted of bark fines, peat moss, pumice, sand, composted manure, and biochar (Lane Potting Mix, Lane Forest Products, Eugene, OR). The initial soil microbiome inoculum was sampled from the top 10 cm of an upland mineral soil under a deciduous forest ecosystem near the University of Oregon campus in Eugene, OR, USA. Each jar was capped and injected with 4.3 mL of 99% CH_4_, which produced a mean headspace concentration of 763.9 ppm (SD = 183.1). Twice per week, jars were flushed in a biosafety cabinet (to avoid contamination) and respiked with CH_4_ to maintain aerobic conditions and elevated CH_4_ concentrations.

For the selection experiment, we created two lines of soil microcosms with 12 jars each: a control line with random selection and an experimental line with directional selection for greater soil CH_4_ oxidation rate. The selection line underwent positive selection where the two or three jars with the highest CH_4_ oxidation rate were homogenized to inoculate the next set of jars. The control line underwent random selection where an equal number of jars as the selection line were chosen at random to inoculate the next set of jars. The number of jars chosen was based on the distribution of fluxes among the positive jars: three jars were chosen in all generations except for passage 3 where two jars were selected. The experiment was carried out over five passages with an average incubation time per generation of four weeks. Methane oxidation rates were determined at the end of the incubation period and selection was performed. For each treatment, the selected jars were homogenized and 5 g of the homogenized soil was used as the living soil inoculum for the next generation. The next set of jars were created in an identical manner to the first generation with fresh autoclaved potting mix and the same moisture and CH_4_ content.

### Methane oxidation rate

Methane oxidation rates were determined after flushing and spiking jars to 1000 ppm CH_4_. Headspace samples of 1 mL were collected from each jar immediately after spiking and then at time points 3, 6, 24, and 48 hours for a 5-point curve. Samples were immediately injected into a SRI model 8610C gas chromatograph equipped with a flame ionization detector (SRI Instruments, Torrance, CA, USA) to determine the headspace CH_4_ concentration. We applied a first-order exponential decay function to determine the rate constant (k, units = d^−1^; i.e., dCH_4_/dt = k[CH_4_]) of the exponential decrease in CH_4_. Oxidation rates are presented as the additive inverse of *k* (i.e., *−k*) so that a more positive value represents a greater oxidation rate. The jars selected to inoculate passage three for the positive selection treatment had the lowest CH_4_ oxidation rate of the twelve jars due to a calculation error in the rate constant.

### Soil DNA extraction and sequencing

A subsample of soil from the starting inoculum and from every jar in passages 2 and 5 was collected and stored at − 80°C. Soil DNA was extracted from 0.25 g soil. Negative controls were extracted from autoclaved potting mix and DNase-free water. Extractions were performed using the DNeasy PowerSoil kit (QIAGEN, Düsseldorf, Germany) and quantified using Qubit dsDNA HS Assay Kit (Thermo Fisher Scientific, Inc., Waltham, MA, USA). To estimate the diversity and relative abundance of the bacterial and archaeal taxa in our soil ecosystems, we sequenced the V4 region of the 16S rRNA gene using the 515F - 806R primer combination (24). PCR mixtures were: 10 *μ*l NEBNext Q5 Hot Start HiFi PCR master mix, 9.2 *μ*l primer mixture (1.09 *μ*M concentration), and 0.8 *μ*l of DNA template. Reaction conditions were: 98°C for 30 s (initialization); 35 cycles of 98°C for 10 s (denaturation), 61°C for 20 s (annealing), and 72°C for 20 s (extension); and 72°C for 2 m (final extension). Reactions were performed in triplicate and then combined. Amplicons were purified twice using 0.8x ratio Mag-Bind RxnPure Plus isolation beads (Omega Bio-Tek, Norcross, GA, USA). Sequencing libraries were prepared using a dual-indexing approach (25,26). Amplicon concentrations were quantified using Qubit and multiplexed at equimolar concentration. Sequencing was performed at the University of Oregon Genomics Core Facility on the Illumina NovaSeq 6000 with paired-end 150 bp reads (Illumina, Inc., San Diego, CA, USA).

### Bioinformatics

Bioinformatics processing was performed in ‘R’ (27). Demultiplexed sequencing reads were denoised using ‘DADA2’ to generate a table of amplicon sequence variants (ASVs) (28). Taxonomic assignment was performed using the Ribosomal Database Project naive Bayesian classifier (29). The presence of contaminants was evaluated using both the prevalence and frequency methods from ‘DECONTAM’ by comparing samples to extraction controls of water (30). Decontam identified 16 potential contaminants based on prevalence and frequency. Visual inspection of abundance-concentration plots indicated that 9 of these were likely contaminants and these ASVs were removed. Amplicon sequence variants that were assigned chloroplast or mitochondria taxonomy were removed prior to analysis.

### Statistical Analysis

Statistical analyses were performed in ‘R’ (27). To determine whether there was a significant change in CH_4_ oxidation rate as a response to selection, we tested a difference in slopes between the selection and control lines. Residuals did not meet the assumptions of constant variance and normal distribution. Therefore, CH_4_ oxidation rates were log_10_ transformed prior to analysis. Following transformation, these assumptions were met. First, we tested if there was a difference of slopes between the selection line and the control based on the interaction between passage and treatment. To test the interaction, we fit two nested models with and without the interaction term and compared them using an F-test with the ‘anova’ function. We then present the slopes for each treatment, which represented the change in CH_4_ oxidation rate per passage as a response to selection.

We estimated the proportion of variation in CH_4_ oxidation rate due to variation in the microbiome as the regression of divergence between the positive line and the control on the cumulative selection differential (31). This estimate is analogous to estimates of “microbiability” from the animal breeding literature, which quantifies the variation in a host trait that is due to microbiome variation (32). The slope of the regression of divergence on cumulative selection differential provides an estimate of realized microbiability (h^2^ ± SE). Divergence was calculated as the mean CH_4_ oxidation rate of the positive treatment minus the mean CH_4_ oxidation rate of the control in each passage. The selection differential was calculated as the difference between the mean of the three selected jars and the mean of all twelve jars in a passage. Cumulative selection differential was calculated as the sum of the selection differential from all preceding selection events. We then regressed cumulative divergence on cumulative selection differential using the ‘lm’ function. We report the slope as percent change by back-calculating the percent change from the log-transformed data into the original units using the formula (10^β^ −1)*100 where *β* is the slope.

Richness was estimated using the method from (33) with a subsample size of 176,545 calculated via the ‘rarefy’ function in ‘vegan’ (34). We tested a difference in richness by both passage and treatment with a Kruskal-Wallace test followed by a pairwise Wilcoxon test. Next, we estimated beta-diversity as the Bray-Curtis dissimilarity by averaging 100 random subsets with a subsample size of 176,545 using the ‘avgdist’ function in ‘vegan’ (34,35). We tested a difference in centroid and dispersion of beta diversity by passage and treatment using a permutational analysis of variance (PERMANOVA) with 999 permutations using the ‘adonis2’ function from ‘vegan’ and tested a difference of group dispersions using ‘betadisper’ and ‘anova’ with 999 permutations (34,36). Lastly, we tested the correlation between CH_4_ oxidation rate and Bray-Curtis dissimilarity in Passage 5 with a distance-based redundancy analysis (dbRDA) using the ‘dbrda’ function in ‘vegan’ and estimated the p-value using a permutation F-test with 999 permutations (34,36)

To identify taxa that responded to selection on CH_4_ oxidation rate, we tested differential abundance between the two treatments in passage 5. We first grouped ASVs at the family level. We chose this level of agglomeration because CH_4_ oxidation is a relatively deeply conserved function (23) and is restricted to a handful of bacterial and archaeal families (37). Therefore, we are most likely to detect an enrichment of methanotrophs at this taxonomic scale. Any ASVs that lacked a family-level taxonomic assignment were grouped at a higher taxonomic level. We then subset the samples in Passage 5 and removed all families with a prevalence of less than 10% in either treatment. We used three methods for testing differential abundance: ANCOM-II, ALDEx2, and CORNCOB (38–41). We then identified the consensus taxa that were significant with all three tests and plotted their relative abundances. For ANCOM-II, we used the ‘ancom’ function in the ‘ANCOM-BC’ package with a cutoff of W = 0.7 (38,39). For ALDEx2, we used the ‘aldex’ function in the ‘ALDEx2’ package with Welch’s t-test and we used an effect size of 1 as our significance threshold (40). Finally, we used CORNCOB with the ‘differentialTest’ function in the ‘corncob’ package with the Wald test and without bootstrapping (41). Lastly, to test differentially abundant methanotrophs, we subset all ASVs within methanotrophic families and tested their differential abundance aggregated at the family and genus level using ‘corncob’. For each test, p-values were adjusted for multiple testing by controlling the false discovery rate using the Banjamini-Hochberg procedure (42).

## Results

### Response to selection on methane oxidation rate

We observed a response to artificial selection on whole-ecosystem soil CH_4_ oxidation rate (Figure 2; difference of slopes: F_2,113_ = 3.85, p = 0.02). At the start of the experiment, the positive selection treatment had a mean CH_4_ oxidation rate that was 24% lower than the control (difference of y-intercepts = -0.34, SE = 0.16, t = -2.14, p = 0.03). There was no change in CH_4_ oxidation rate in the control over the five passages (slope = -0.01, SE = 0.05, t = -0.26, p = 0.80). By contrast, the selection treatment had a 50.7% increase in CH_4_ oxidation rate per passage (slope = 0.18, SE = 0.06, t = 2.76, p = 0.01).

**Figure 2:**
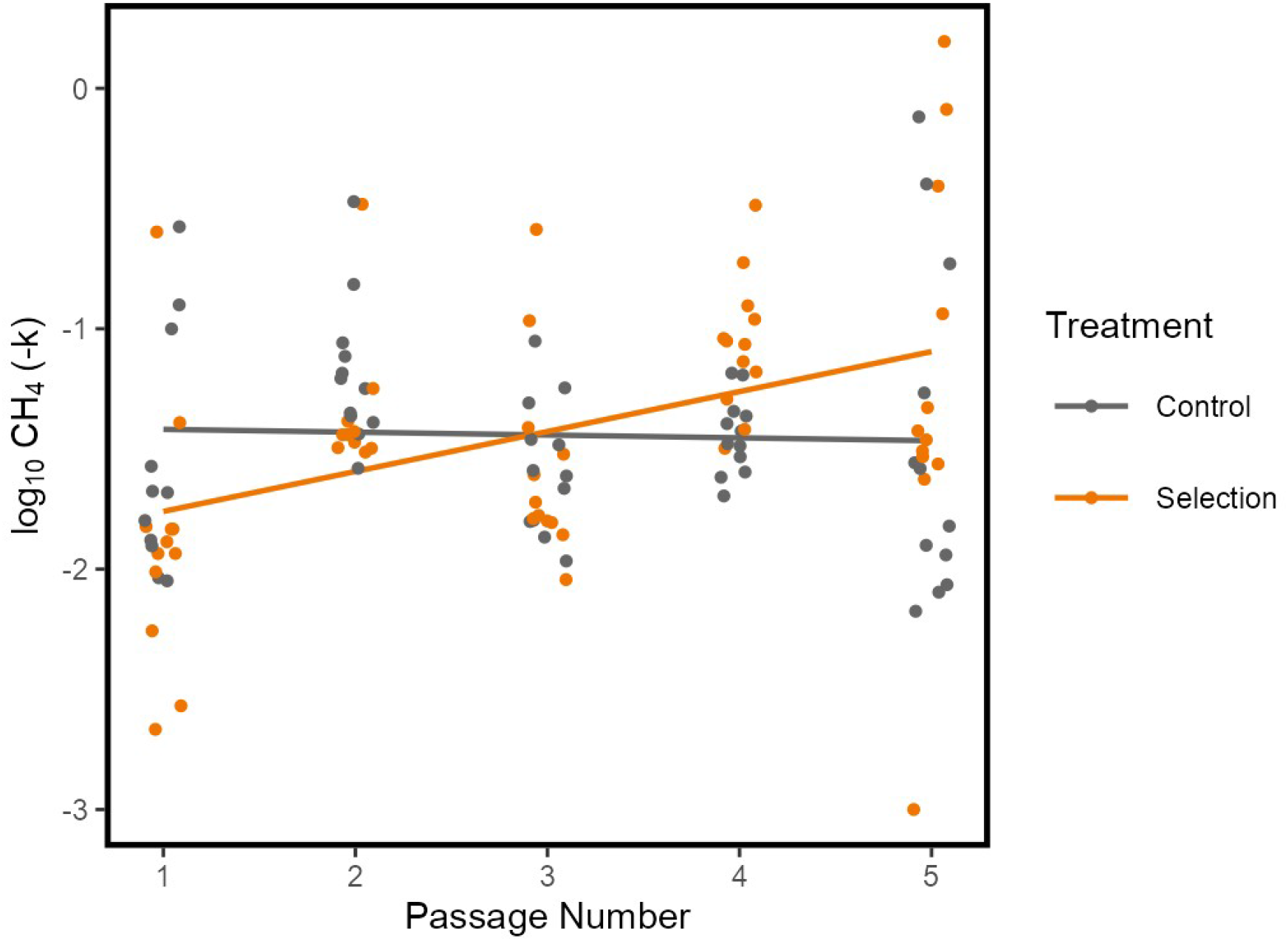
Response to selection on soil CH_4_ oxidation rate. The y-axis is CH_4_ oxidation rate as the log_10_ of the additive inverse of the first-order exponential decay constant *k* (i.e., *−k*) with units day^-1^ so that a more positive value represents a higher CH_4_ oxidation rate. Orange points and regression line are for the positive selection treatment and gray points and regression line are for the control. There was a significant difference of slopes between the positive selection treatment and the control (F_2,113_ = 3.85, p = 0.02).

To estimate the proportion of variation in CH_4_ oxidation rate due to variation in microbiome composition–i.e., microbiability (as described in the Methods; (32)–we regressed divergence between the positive selection treatment and the control against the cumulative selection differential. The microbiability was 0.31 ± 0.17, though this was not significant (F_1,2_ = 3.44, p = 0.20).

### Taxonomic richness

Median ASV richness decreased from 3406.6 (778.5) in passage 2 to 1557.8 (157.7) in passage 5 (Kruskal-Wallace test: χ^2^ = 35.4, df = 3, p < 0.001; pairwise Wilcoxon test: p < 0.001). However, there was no difference in richness between the selection treatment and the control in passage 2 or 5 (pairwise Wilcoxon test; Passage 2: p = 0.66, Passage 5: p = 0.67). In addition, there was no correlation between richness and CH_4_ oxidation rate across the two treatments in passage 5 (Spearman’s rho = -0.2, p = 0.3).

### Community dissimilarity

Bray-Curtis dissimilarity of the soil microbiome varied strongly by passage and weakly by treatment with an interaction between passage and treatment (Figure 3). Passage explained 55.9% of the variation in Bray-Curtis dissimilarity (F_1,44_ = 73.3, p = 0.001), treatment explained 5.9% of the variation (F_1,44_ = 7.8, p = 0.001), and the interaction between treatment and passage explained 4.7% of the variation (F_1,44_ = 6.2, p = 0.003). There was no difference in dispersion between treatments or passages (F_3,44_ = 0.91, p = 0.45). Finally, CH_4_ oxidation rate was correlated with Bray-Curtis dissimilarity across both treatments in passage 5 and explained 9.6% of the variation in Bray-Curtis dissimilarity (dbRDA: F_1,22_ = 2.34, p = 0.010)

**Figure 3:**
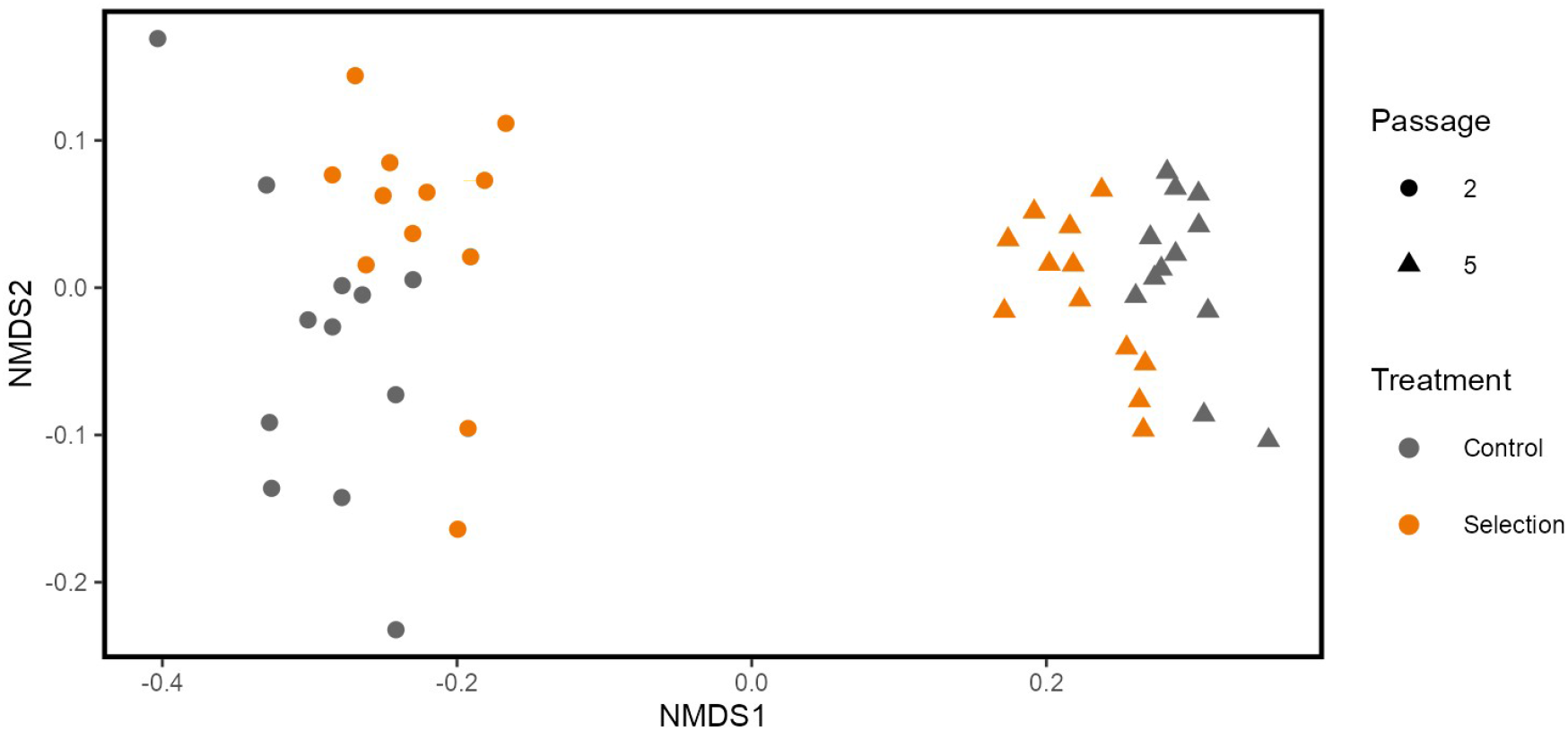
Non-metric multidimensional scaling plot of beta diversity for all jars. Dissimilarities are rarefied Bray-Curtis dissimilarity averaged over 100 subsamples. Orange points are the positive selection treatment and gray points are the control. Circles are passage 2 and triangles are passage 5.

### Taxa that responded to selection

To identify taxa that responded to selection on soil CH_4_ oxidation rate, we tested the differential relative abundance of families in the selected jars relative to the control jars within passage 5 using three methods and then plotted the taxa identified by all three methods. We identified 12 families that were enriched or depleted in the selection treatment relative to the control (Figure 4).

**Figure 4:**
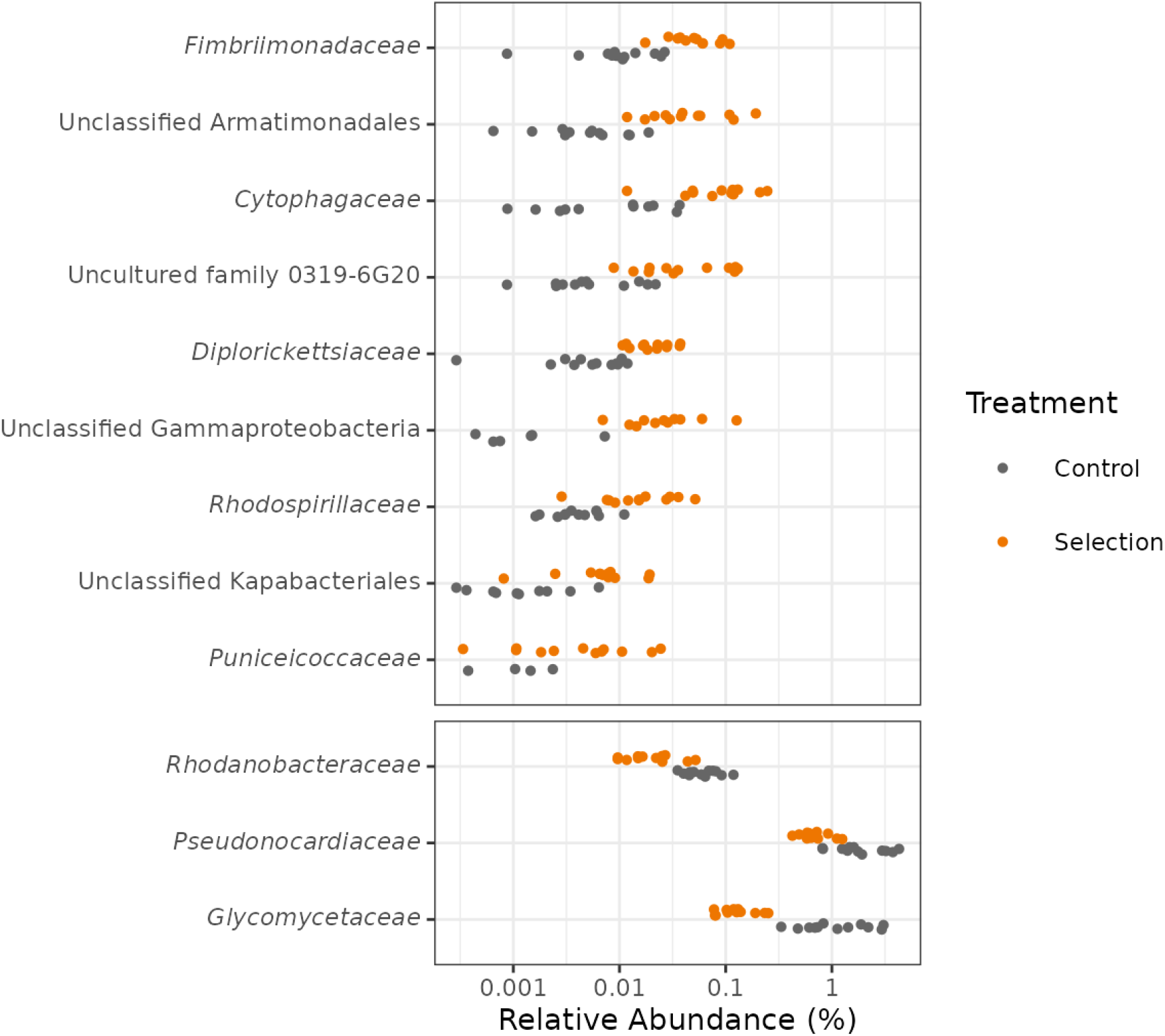
Differentially abundant family-level taxa identified by ANCOM-II, ALDEx2, and CORNCOB. Values on the x-axis are relative abundances on a log_10_ scale. Taxa in the top panel are enriched in the positive selection treatment relative to the control and taxa in the bottom panel are depleted in the positive selection treatment relative to the control. Taxa are sorted by their effect size with taxa at the top having the largest positive effect size and taxa at the bottom with the largest negative effect size.

Overall, none of the families enriched in the selection treatment contain known methanotrophs. Several taxa identified had a higher taxonomic designation that contains methanotrophs, for example, the Gammaproteobacteria class had a large effect size. The Gammaproteobacteria include the type I and type X methanotrophs in the families *Methylococcaceae* and *Methylothermaceae* (43). However, the Gammaproteobacteria is among the most diverse groups in the Prokaryotes, so this is not strong evidence for a selection response by methanotrophs (44). In addition, the *Puniceicoccaceae* is a member of the phylum Verrucomicrobia. The Verrucomicrobia is a diverse group that contain known methanotrophs as well as ammonia-oxidizing bacteria (45). Other than these two groups, none of the other taxa enriched in the selection treatment are known to be related to methanotrophs. Two groups in the Armatimonadales were enriched in the selection treatment including the family *Fimbriimonadaceae* and an unclassified ASV from the order Armatimonadales (46). *Cytophagaceae* was also enriched in the selection treatment and contains a number of mainly aerobic heterotrophs that can digest a variety of macromolecules (47). The remaining families include the uncultured family 0319-6G20, *Diplorickettsiaceae*, *Rhodospirillaceae*, and an unclassified Kapabacteriales.

We did not identify any methanotrophic families in the overall differential abundance analysis. However, we wanted to look more closely at the known methanotrophs in our dataset to be sure that they did not have an effect. To do this, we subset the ASVs in our dataset that were in families that contained methanotrophs. Only two families were represented: *Methylacidiphilaceae* and *Beijerinckiaceae*. Aggregating reads at the family level, neither family was differentially abundant between the two treatments. However, aggregated at the genus level, a group of unclassified genera in the *Beijerinckiaceae* were depleted in the selection treatment and the genus *Rhodoblastus*, a member of the *Beijerinckiaceae*, was enriched in the selection treatment. While many *Beijerinckiaceae* are methanotrophs, several taxa in this family have lost the ability to oxidize CH_4_ and it appears that *Rhodoblastus* species are not able to grow on CH_4_, though they can grow on methanol (48,49). Based on this analysis, it appears that no methanotrophs were enriched in the positive selection treatment.

## Discussion

We used artificial ecosystem selection to estimate the contribution of variation in microbiome composition to variation in the rate of an ecosystem function, CH_4_ oxidation in soil, independent of environmental variation. Understanding how and to what degree microbiome variation contributes to variation in ecosystem function is important for many reasons. For example, successful microbiome manipulations require that the manipulated microbiome contribute to variation in ecosystem function independent of other drivers of ecosystem variation (such as variation in environmental conditions). This is because the drivers of variation in ecosystem functions can interact in complicated ways (Figure 1); for example, environmental variation can indirectly contribute by providing conditions that select for microbial groups that in turn alter the rates of ecosystem functions. Determining the independent contribution of microbiome variation to ecosystem function is crucial because if microbiome composition is driven primarily by environmental conditions, then introducing a desirable taxon through microbiome manipulation without altering the environment will likely be unsuccessful at shifting the targeted ecosystem function. The artificial selection approach is different from the comparative and manipulative approaches used in past attempts at answering this question, because it can control for both the direct effect of environment on function as well as the indirect effect on environment via its impact on microbiome assembly.

In our study, we observed an increase in CH_4_ oxidation rate in the selection treatment relative to the control, which demonstrates that there was a response to selection. Given that we observed a response to selection, we conclude that variation in the microbiome contributes to variation in the CH_4_ oxidation rate independent of the environment. This suggests that microbiome manipulations could be an effective approach for altering the rate of CH_4_ oxidation in this soil, and that the artificial selection approach may be useful in determining the potential for microbiome manipulations for other functions in other ecosystems.

Given that variation in the microbiome is associated with variation in the rate of an ecosystem function in our system, a reasonable follow-up question is “how much variation in ecosystem function is associated with microbiome variation in this system?” One way to estimate this is to determine how much the recipient jars resemble the selected donor jars that were used to inoculate them (31). We can calculate the response to selection as the difference between two successive passages in their mean CH_4_ oxidation rate. We will denote this as *R*. We can also calculate the strength of selection as the difference in mean CH_4_ oxidation rate between the twelve jars in one generation and the three jars chosen for selection in that generation, which we will call the selection differential and denote as *S*. If we plot the cumulative *R* against the cumulative *S*, the slope of this relationship will equal the proportion of variation explained by the microbiome. If the change in mean function from passage one to passage two (*R*) is equal to the difference in mean function between the twelve jars in passage one and the three jars selected to inoculate passage two (*S*), then we would conclude that 100% of the variation is due to variation in the microbiome. Likewise, if recipients do not resemble the donors in their mean CH_4_ oxidation rate and simply wander randomly, then we would conclude that all of the variation is due to the environment or technical variation.

The relationship between microbiome variation and ecosystem function variation is analogous to the concept of “heritability” (50) used by quantitative geneticists, or more precisely the concept of “microbiability” (32) proposed by microbiome scientists who study host-associated microbiomes. Although rarely used in the study of environmental microbiomes, this concept could be very useful for understanding and manipulating microbially-mediated functions in a variety of ecosystems. In our experiment, variation in microbiome taxonomic composition statistically explained (i.e., was associated with) 31.5% of the variation we observed in the rate of CH_4_ oxidation, though this was not significant. However, we did observe a significant divergence between the positive selection and control lines, which suggests that the imposed selection and passaging of microbiomes was sufficient to generate variation in soil CH_4_ oxidation rate. Future studies with greater replication could more precisely estimate the microbiability. This suggests that there is substantial potential for altering this ecosystem function through microbiome manipulation in this soil. It is very likely that the “environmental microbiability” will be different for other ecosystem functions in this soil and for CH_4_ oxidation in other soils. However, our experiment demonstrates that this relationship is measurable and provides an example of how this can be accomplished.

We next wanted to determine which aspects of the microbiome might explain the divergence in CH_4_ oxidation rate between the two treatments. There are three inter-related ways that microbiomes could have responded to selection in this experiment: gain or loss of taxa, changes in the relative abundances of taxa, or changes within the genomes of the constituent taxa. We surveyed microbiome variation via 16s rRNA ribotyping in our experiment, which allowed us to deeply sample taxonomic diversity but did not allow us to directly address whether taxa in this experiment evolved genomic changes as a result of selection. However, if such genomic changes resulted in increased persistence or abundance of the population with these changes, this would be detectable. Therefore, we will focus on the first two possibilities.

Richness at the ASV level did not vary between the two treatments and there were relatively few taxa gained or lost in the selection treatment and none of these were prevalent across the 12 jars in passage 5. Therefore, the gain or loss of species is unlikely to explain the increase in CH_4_ oxidation rate. However, we found that Bray-Curtis dissimilarity was greater between the two treatments in Passage 5 than within each treatment and was correlated with CH_4_ oxidation rate, which suggests that changes in the relative abundance of taxa could explain the response to selection.

Even though we observed an increase in CH_4_ oxidation rate in the selection treatment and a difference in composition between the two treatments, we did not observe an increase in the relative abundance of known methanotrophs. This was surprising given that CH_4_ consumption is not a common trait among microbes and that it is often assumed that the rate of an ecosystem function is limited by the final enzymatic step in the underlying metabolic pathway (11). In certain ecosystems, CH_4_ production and consumption are correlated with the abundance of methanogens and methanotrophs as estimated from marker genes (18,19). However, our results suggest that in this system ecosystem-scale CH_4_ oxidation rates can be altered by non-methanotrophs, perhaps through ecological interactions with methanotrophic species, or by unknown methanotrophs. This suggests that simple assumptions about how microbes contribute to rate variation in ecosystem function may not apply universally, and it demonstrates the importance of using biologically “agnostic” approaches (that make few starting assumptions) to linking microbial taxa to ecosystem functions (51). Artificial ecosystem selection is an important example of such an approach.

There is increasing interest in using artificial selection for understanding and manipulating the microbiomes associated with plants and animals (a.k.a., “microbiome breeding”; (52)). Our study demonstrates that artificial ecosystem selection can also be an important tool for exploring the relationship between microbiome composition and ecosystem function in non-host systems. This approach can provide unique information about the independent contribution of microbiomes to ecosystem functions. Such information is crucial if we are to successfully manipulate environmental microbiomes to alter ecosystem functions, whether to improve crop productivity (53) or ameliorate the impacts of environmental change (54).

## Acknowledgments

This project was supported by the National Science Foundation Graduate Research Fellowship Program (grant no. DGE 1255832) and the ARCS Foundation Florence and Mike Nudelman Scholarship. Figure 1 was created with BioRender.com.

## Data Availability Statement

The 16S rRNA sequencing data generated during the current study are available in the NCBI Sequence Read Archive (SRA) under BioProject accession number PRJNA832314, https://www.ncbi.nlm.nih.gov/sra/PRJNA832314. The metadata generated during the current study as well as the scripts to recreate the analysis are available on Github, https://github.com/amorris28/artificial_ecosystem_selection.

